# Detecting Non-linear Dependence through Genome Wide Analysis

**DOI:** 10.1101/2025.02.12.637804

**Authors:** Wonuola A Akingbuwa, Michel G Nivard

**Affiliations:** Department of Biological Psychology, Vrije Universiteit Amsterdam, Amsterdam, the Netherlands; Amsterdam Public Health Research Institute, Amsterdam, the Netherlands; Medical Research Council Integrative Epidemiology Unit, University of Bristol, Bristol, United Kingdom; Population Health Sciences, Bristol Medical School, University of Bristol, Bristol, United Kingdom

## Abstract

In the current study we introduce statistical methods based on trigonometry, to infer the shape of a (non)-linear bivariate genetic relationship. We do this based on a series of piecemeal GWASs of segments of a target (continuous) trait distribution, and the genetic correlations between those GWASs and a second trait. Simulations confirm that we are able to retrieve the shape of the relationship given certain assumptions about the nature of the relationship between the traits. We applied the method to the genetic relationship between BMI, sleep duration, and height, and psychiatric disorders (ADHD, anorexia nervosa, and depression) using data from approximately 450K individuals from UK Biobank.

In the relationship between BMI and psychiatric traits, we found that the expected value of depression is a nonlinear function of BMI i.e. there is a nonlinear genetic relationship between both traits. We observed similar findings for the genetic relationship between BMI and anorexia, sleep duration and depression, and sleep duration and ADHD. We observed no underlying nonlinearity in the genetic relationship between height and psychiatric traits.

Using a novel statistical approach, we show that nonlinear genetic relationships between traits are detectable and genetic associations as quantified using global estimators like genetic correlations are not informative about underlying complexities in these relationships. Our findings challenge assumptions of linearity in genetic epidemiology and suggest that bivariate genetic associations are not uniform across the phenotypic spectrum, which may have implications for the development of targeted interventions.

## INTRODUCTION

In genetic analyses nonlinearity is primarily invoked in the relationship between genotypes and traits. In these types of studies nonlinearity is defined as a dominance effect - where one allele masks the effects of another variant at the same locus, epistasis - where the effect of one variant is modified by another, or gene-environment interaction - differential responses to environmental variation based on genotype differences ^1–3^. However another potential form of nonlinearity can arise from nonlinearity in the relationship between pairs of heritable traits. Famous examples are U-shaped, or J-shaped associations, where both low and high levels of an exposure are associated with an outcome. Currently, nonlinearity in the bivariate genetic relationship between phenotypes is generally left unmodelled. While estimators like the genetic correlation do not stipulate the relationship between traits is linear, the estimate is entirely uninformative about the nature or shape of that relationship. We introduce two estimators - based on polynomial or cubic spline approximation - of the shape of the relationship between the genetic liabilities of two traits, using GWAS summary statistics.

Non-linear, and other complex phenotypic associations between traits are generally well documented/explored. There is evidence that in at least a subset of people, income is positively associated with subjective well-being up to a certain point, after which the effect diminishes or plateaus ^4,5^. Non-linear dose-response relationships have also been reported between physical activity and health outcomes; moderate, rather than low or high exercise intensity is associated with mood improvements ^6^, and significant reduction in all-cause mortality with increasing physical activity up to a threshold, after which the effect flattens out ^7,8^. Finally, a J- or U-shaped association has been observed between body mass index (BMI) and all-cause mortality such that both high and low BMI are associated with increased risk of mortality ^9,10^. It is important to acknowledge that these associations are not evidence of causal relationships between these phenotypes, and some of these associations are affected by confounding. For example, the J-shaped association between alcohol consumption and all-cause mortality which suggests a protective effect of light or moderate alcohol use compared to abstainers has been shown to disappear when controlling for various sources of bias ^11,12^. However, regardless of causality, it is important to be able to establish that these nonlinear associations exist in the first place in order to study them.

In the case of psychiatric traits, nonlinear relationships have been observed between BMI and depression, as well as between sleep duration and mental health. Both short and long sleep duration are associated with anxiety, depression, and mania symptoms, as well as self-harm behaviours and negative affect ^13–15^. Similarly, U- or J-shaped associations have been observed for the relationship between BMI and depression, such that both high and low BMI are associated with increased depression ^16,17^. This relationship is characterised by a negative association initially, and then a positive association. Nevertheless, a decent proportion of the research on this association has examined the linear relationship with contradictory results; findings range from positive to negative, to null/insignificant ^17^.

Prior rudimentary nonlinear genetic of analysis of sleep, where the phenotype “how many hours do you sleep on average” was split into 3 traits: total sleep duration, sleeping longer than average (versus sleeping an average amount), and sleeping less than average (versus sleeping an average amount) clearly highlights the need for nonlinear genetic analysis ^18^. Both sleeping longer and sleeping less than average are significantly genetically related to depression symptoms and various other adverse health outcomes and traits, while total sleep duration is not genetically related to depression symptoms (and other traits), likely due to those effects cancelling out in aggregate. The genetic association as quantified by the genetic correlation coefficient between total sleep duration and other traits, were not informative about the nature of the relationship between both sets of traits; they are not informative about any phenotypic heterogeneity that might exist and any resulting nonlinearity is not reflected in estimates of these associations and can go undetected entirely. In such analyses or estimates, the entire spectrum of the trait is generally assumed (or treated) as having the same underlying genetic liability, and fails to account for the possibility of differentiation of genetic architecture across the (continuous) phenotype spectrum.

In order to fully model nonlinearity in cross-trait genetic associations, we introduce a method (and accompanying R package, “TriGenometry”) to interpolate the shape of a (non)-linear relationship between two traits based on a series of piecemeal GWASs of segments of the target (continuous) trait distribution, and the genetic correlations between those GWASs and a second trait. We then estimate the genetic dependence between pairs of traits while imposing minimal prior constraint on the shape of the dependence. The presence of nonlinearity might indicate that there is differentiation of genetic risk across the phenotype spectrum in the genetic relationship between two variables. We test the method on BMI, sleep duration, and height, investigating nonlinear relationships with depression, ADHD, and anorexia nervosa.

## METHODS

### Conceptual model

Assuming two phenotypes (e.g. BMI and depression) to be the sum of additive genetic (*a*_*bim*_ and *a*_*dep*_) and environmental effects (*e*_*bim*_ and *e*_*dep*_), we are interested in estimating the functional relationship that describes the expected value of *a*_*dep*_ given a value of *a*_*bim*_. We estimate the function *f*(*a*_*bim*_) from multiple estimates of the derivative *f*′(*a*_*bim*_), each based on genome-wide association study (GWAS) analyses of pairs of sections of BMI, in a procedure that is described in more detail below.

### Statistical method

#### Estimating nonlinear relationship

For the sake of simplicity, we consider traits *x* and *y* as the sum of additive genetic (*a*_*x*_ and *a*_*y*_) and environmental (*e*_*x*_ and *e*_*y*_) effects as follows:

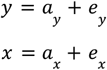

We were interested in estimating the functional relationship that describes the expected value of *a*_*y*_ given a value of *a*_*x*_ :

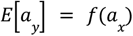

Where *y* can be binary, ordinal, or continuous and *x* must be continuous (or practically so) to facilitate estimating the continuous function: *f*(*a*_*x*_). In the current analyses *y* are psychiatric disorders (ADHD, anorexia nervosa and depression), and for *x* we considered BMI, height, and sleep duration. Note that unlike a linear relationship, [*E a*_*y*_] = *f*(*a*_*x*_) and the inverse *E* [*a*] = *f*(*a*_*y*_) are not simple transformations of each other, and the particular direction of the equation should not be interpreted as a commitment to a causal direction, or causal nature of the relationship. We learn how the expectation of *y* depends nonlinearly on *x*, which does not give us full information of the expectation of *x* given *y*.

We estimate the function *f*(*a*_*x*_) based on multiple estimates of the distance: *f*′(*a*_*x*=*i*_) − *f*′(*a*_*x*=*j*_) between pairs of values (*i* and *j*) along the continuum of *x*. Using the example of BMI, we do this by binning BMI (*x*) into 30 quantile-based bins (ensuring sufficient sample size in each bin), and performing a GWAS of all pairwise combinations of bins.

For each SNP in the HapMap 3 reference set we estimate:

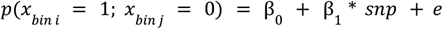

Where an individual’s value for *x* (BMI) is either in *bin i*, or in *bin j*, such that the group of individuals with BMI values in each bin are “assigned” to that bin and serve as either the case or control group in the series of GWAS contrasting each pair of bins. Based on each of these GWASs, which characterise cross sections of *x*, we then estimate the genetic correlation with an external trait *y*. As both the distance between the bins, *dx*, and the genetic correlation between the GWAS of that distance with trait *y* are known, we use the fact that a correlation can be transformed to an angle, to estimate the unknown distance on *y* given the distance on *x*. By distance *dx* we refer to the difference between the mean of estimates in *x* _*bin i*_ and *x*_*bin j*_ on a reasonable scale (usually the original scale of *x*). In the current analyses we primarily use the median value of *x* in each bin.

First we divide the genetic correlation estimates (*cor*) by the distance between each pair of bins (*dx*), and transform it to the acute angle of a right triangle:

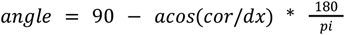

Then we can calculate the estimated distance on *y* (*dy*) as it is the opposing side of a triangle with an adjacent side with length *dx* and a known acute angle (Figure 1):

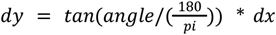

**Figure 1:**
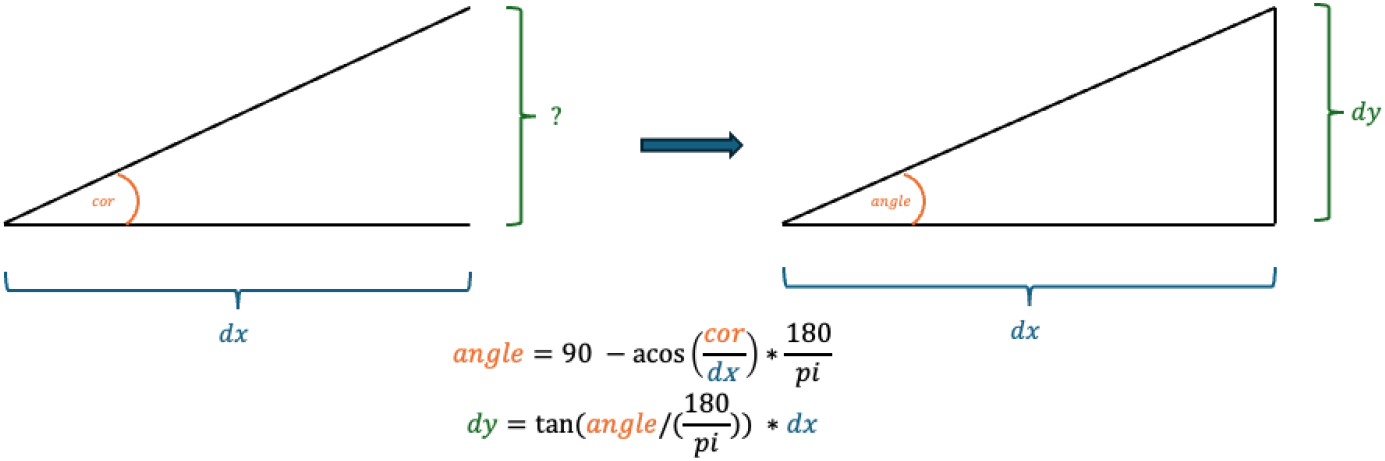
Visualisation of conversion from correlation to angle

Distances *dy* are all relative to a specific pair of bins of *x*, where *dx* is the distance between the midpoints of the bins. Given 30 (BMI) bins, we perform 435 GWASs between bin pairs, which translate to 435 estimated distances on *y*.

#### Inferring the function from the set of paired distances

We have now estimated distances on 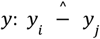 that correspond to known distances *x*_*i*_ − *x*_*j*_ and based on these we want to estimate the functional relation:

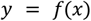

We can conveniently write:

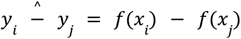

And then use a cubic spline function (alternatively a 5th degree polynomial) of *x*, which best fits the observed set of distances:

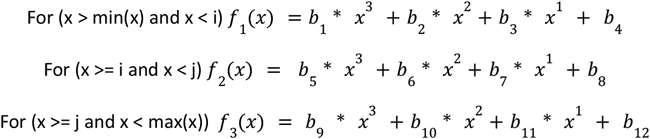

With the following constraint on the equations introduced during optimisation:

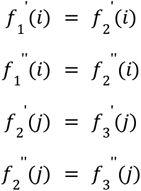

The parameters of which are estimated through numerical optimization, minimising the following function:

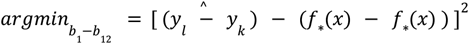

Where * indicates functions 1, 2, or 3 are substituted in, dependent on the value of *x*. The cubic-spline we consider consists of 3 local 3rd degree polynomials that are fit to segments of the data. The 3 polynomials are optimised with the constraint that neighbouring polynomials have identical 1st and 2nd derivatives where they meet (i.e. collectively produces a smooth function). Alternatively the cubic spline approximation can be substituted with a polynomial function. We further validate the ability of our method to retrieve nonlinear relations using simulations (Supplemental Text; Supplementary Figure 1). Code for TriGenometry can be found here: https://github.com/WonuAkingbuwa/trigenometry.

#### Assumptions

We bin the observed trait BMI, or *x*, not the additive genetic component *a*_*x*_ and so an assumption we return to in simulation is that binning *x* is sufficiently informative of *a*_*x*_. We do not observe *f*′(*a*_*x*=*i*_) − *f*′(*a*_*x*=*j*_), rather we perform a GWAS of *f*(*x*) − *f*(*x*).

If all genetic effects on *x* are fully linear and additive, then GWAS of individuals binned on variable *x* are expected to be genetically perfectly correlated. In order for a nonlinear relationship between *x* and *y* to be detected we require at least some genetic effects on *x* to be heterogeneous across *x*. This certainly occurs if heritable trait *y* has a heterogeneous influence across values of *x* (e.g. when *y* can cause *x* to go up or down) as effects on *y* would heterogeneously impact *x*. Alternatively, if through other processes variants have heterogeneous effects on *x* and *x* has a nonlinear effect on *y* the nonlinear relationship could be detected. However, if all genetic effects on *x* are additive and homogeneous across *x* (i.e absolutely no GxE and no GxE that results in the genetic effect differing across levels of *x*), a nonlinear relationship where *x* is causal for *y* can go undetected by our method (see simulated violation of assumption in Supplementary Text; Supplementary Figure 2).

#### Assessment of uncertainty

There are various sources of uncertainty in our procedure; first the genetic correlations are estimated with variable precision. Second, approximation of a function, whether by polynomial or cubic spline, introduces further uncertainty. In order to assess the uncertainty around the resulting curve from estimating the function of *x*, we perform 100 random resamplings of the genetic correlations taking into account their standard errors and redraw the curve 100 times.

### Empirical analysis

#### Sample

Participants were from the UK Biobank and data was obtained under project ID 40310. The UK Biobank is a population-based cohort of over 500,000 individuals with genotype and phenotype data from the United Kingdom. Participants were aged between 40 and 69 years old at recruitment ^19^.

#### Genetic analyses

##### Genotyping and quality control

Genotypes were imputed to the Haplotype Reference Consortium (HRC) reference panel.

Pre-imputation quality control (QC) and imputation have been previously described ^19^. We extracted single nucleotide polymorphisms (SNPs) from the HapMap3 CEU reference panel (1,345,801 SNPs) from the imputed dataset. We need only these SNPs (as opposed to full genotypes) because subsequent genetic correlations are estimated using LD score regression which uses the HapMap3 reference panel to estimate LD and drops all SNPs not present in the reference file.

Pre-principal-component analysis (PCA) QC was performed on unrelated individuals, filtering out SNPs with minor allele frequency MAF < 0.01 and missingness > 0.05, which left 1,252,123 SNPs. Individuals with non-European ancestry were subsequently filtered out, and SNP QC was repeated for unrelated Europeans (N = 312,927), filtering out SNPs with MAF < 0.01, missingness > 0.05 and HWE p < 10^-10^, leaving 1,246,531 SNPs. The HWE p-value threshold of 10^-10^ was based on: http://www.nealelab.is/blog/2019/9/17/genotyped-snps-in-uk-biobank-failing-hardy-weinberg-equilibrium-test. As a result genotype data was available for 456,028 individuals with European ancestry, with 1,246,531 QC-ed SNPs remaining in the final dataset. Ancestry was determined using PCA in GCTA and has been previously described ^20^.

##### BMI GWAS

BMI information was obtained from UKB data-field 21001 “Body mass index (BMI)”. BMI was calculated by dividing participants’ weight in kilograms by the square of their height in metres. Phenotype and genotype information was available for 454,441 participants. Participants were stratified into one of 30 quantile-based “bins” according to their BMI. Bins were smaller at the extremes of the BMI distribution and became larger with increasing proximity to the middle of the BMI distribution, and were chosen such that they emphasise the tails of the distribution as we hypothesise that that is where specific genetic signal might show up. From low to high BMI, participants were stratified into 2 half percentile bins (e.g. 0%-0.5%), 4 one percentile bins (e.g. 2%-3%), 18 five percentile bins (e.g. 10%-15%), 4 one percentile bins (e.g. 95-96%), and 2 half percentile bins (99%-99.5%), depending on the position of their BMI in the total sample distribution.

We performed case-control linear mixed model association testing analyses of the BMI bins using fastGWA implemented in Genome-wide Complex Trait Analysis (GCTA) ^21,22^. The GWASs were made up of all 435 pairwise combinations of all BMI bins, with the higher BMI bin serving as the control group in each GWAS instance. Analyses were corrected for age, sex, 25 PCs, and genotyping batch. Total GWAS sample sizes ranged from 4373 to 45,859 individuals depending on the percentile pairs, with case sample size ranging from 2177 to 22951 (Supplementary Table 1). Where the genetic variance estimate was not significant, GCTA used linear regression for the association tests.

##### Sex-stratified analyses

Participants were also stratified by sex. For each sex, we repeated the phenotype stratification process separately and performed GWAS analyses as described above in the combined sample. Total sample sizes from the female GWASs ranged from 2324 to 25095, while they ranged from 1984 to 20977 for the male GWASs (Supplementary Table 1).

##### Genetic correlations

Genetic correlations between BMI GWASs and psychiatric traits were estimated using LD score regression (LDSC) ^23^. LDSC employs a metric called LD score, which is computed for each SNP by summing the correlations between that SNP and all nearby SNPs. This calculation is performed using an ancestrally similar reference sample. The genome-wide LD information used was based on European populations from the HapMap3 reference panel. Genetic correlations are estimates of the degree of shared variance between traits that is attributed to all the measured SNPs. They are estimated from the resulting slope of the regression of the product of the two z-scores from the two GWASs on the LD score. Genetic correlations with psychiatric disorders were based on recent GWAS of anorexia nervosa ^24^, ADHD ^25^, and depression ^26^.

##### Sleep duration

Information on sleep duration was obtained from UKB data-field 1160 “About how many hours sleep do you get in every 24 hours? (please include naps)”. We excluded participants who responded “Prefer not to answer” or “Do not know”. Phenotype and genotype information was available for 453,130 participants. Participants were stratified into one of 9 bins based on number of hours slept; <4 hours, 4, 5, 6, 7, 8, 9, 10, and >10 hours. We performed 36 pairwise case-control linear mixed model association testing analyses of the 9 sleep duration bins in the same manner as we did for BMI. Total GWAS sample sizes ranged from 2637 to 309,442, with case sample size ranging from 826 to 176,764 (Supplementary Table 2). Genetic correlations with psychiatric traits were estimated using LDSC.

##### Height

Height was included as a “negative control” as there was no expectation of a nonlinear association with any of the psychiatric traits. Participants’ heights in cm were obtained from UKB data-field 50 “Standing Height”. Phenotype and genotype information was available for 454,932 participants, who were subsequently stratified into 30 percentile bins as was done for BMI. All subsequent analyses matched what was done for BMI, although we did not perform sex stratified analyses. GWAS sample sizes ranged from 2640 to 68,456 with case sample size ranging from 886 to 35575 (Supplementary Table 3). Genetic correlations were estimated using LDSC.

## RESULTS

### BMI

Analyses for which genetic correlations could not be estimated by LDSC were excluded; all estimates were included in the nonlinear function regardless of significance, although out of bound estimates (> ±1) are also excluded as TriGenometry limits estimates between −1 and 1. Full LDSC results from the binned GWAS analyses, including sex-stratified analyses, are provided in Supplementary Tables 4-6 and Supplementary Figures 3-5.

Significant aggregate genetic correlation estimates have been reported for depression (r_g_ = 0.08, SE = 0.023, P = 0.006) ^26^, ADHD (r_g_ = 0.26, SE = 0.032, P = 1.68E-15) ^25^, and anorexia (r_g_ = −0.32, SE = 0.03, P = 8.93E-25) ^24^. After applying the nonlinear function to the genetic correlation estimates, we observe a nonlinear genetic relationship between BMI and depression similar to the observed relationship.

Specifically, the expected value of *a*_*dep*_ (*E*[*a*_*dep*_]) is a nonlinear function of *a*_*bim*_ (*E*[*a*_*bim*_]) and both high and low BMI are genetically associated with depression. On the other hand, the association between ADHD and BMI is less clear as both nonlinear functions do not converge on a clear shape (Figure 2). Contrary to negative linear correlation suggested by global estimates of the association between anorexia nervosa and BMI, we find that this finding is confined to BMI <30 and then plateaus at higher BMI in males, suggesting low to no genetic correlations between anorexia and high BMI. This association is more pronounced in females, and in females with BMI > 30, there is some indication that the relationship between anorexia and BMI changes from negative to positive (Figure 3, Supplementary Figure 6). Results of sex-stratified analyses for ADHD and depression are provided in Supplementary Figures 7 and 8.

**Figure 2:**
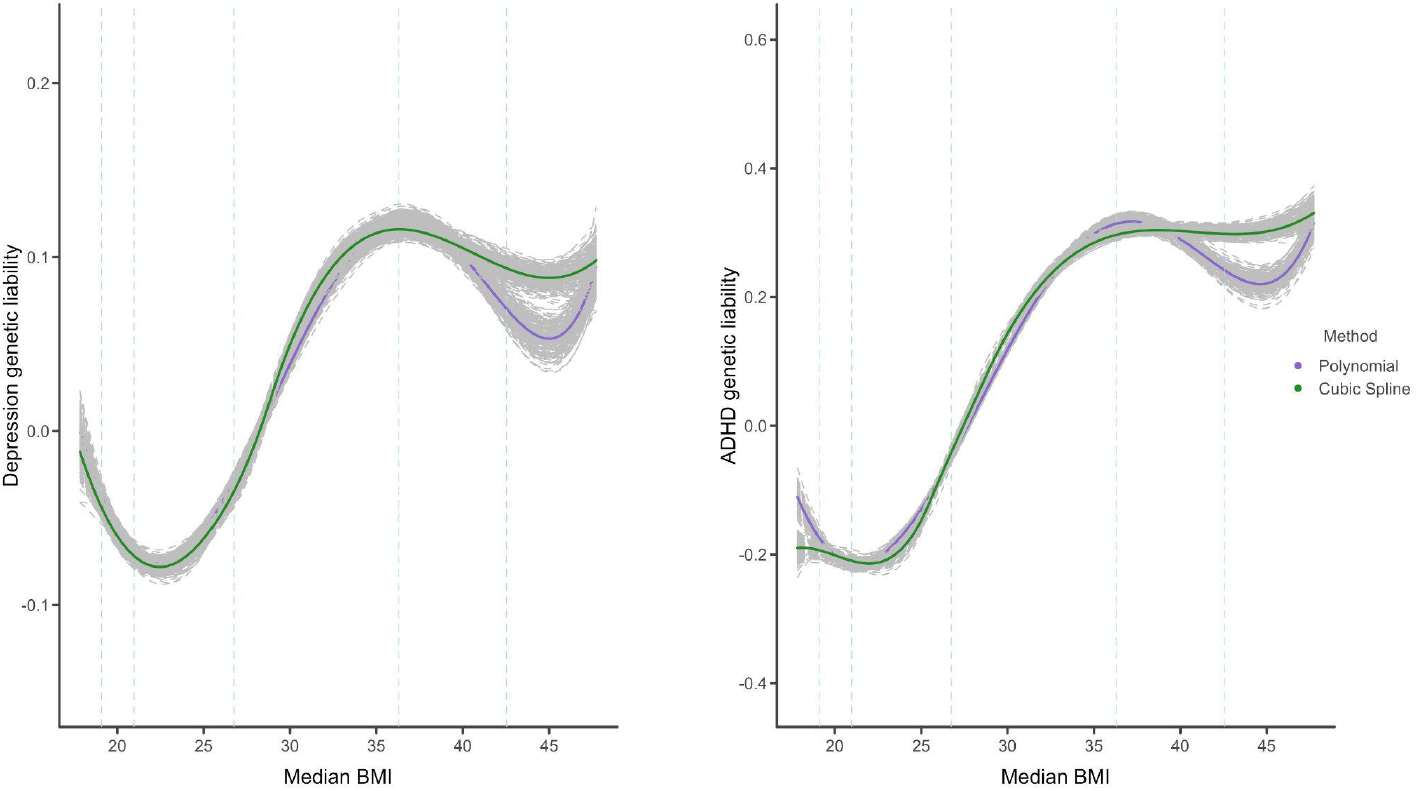
Genetic relationship between BMI and depression/ADHD ***Note***: median BMI per bin is plotted on the *x-axis and the y-axis represents the expected liability of depression/ADHD given BMI on a relative scale (scale values cannot be compared between outcomes). Dashed lines represent BMI values at the 1st, 5th, 50th, 95th and 99th percentile of the data*

**Figure 3:**
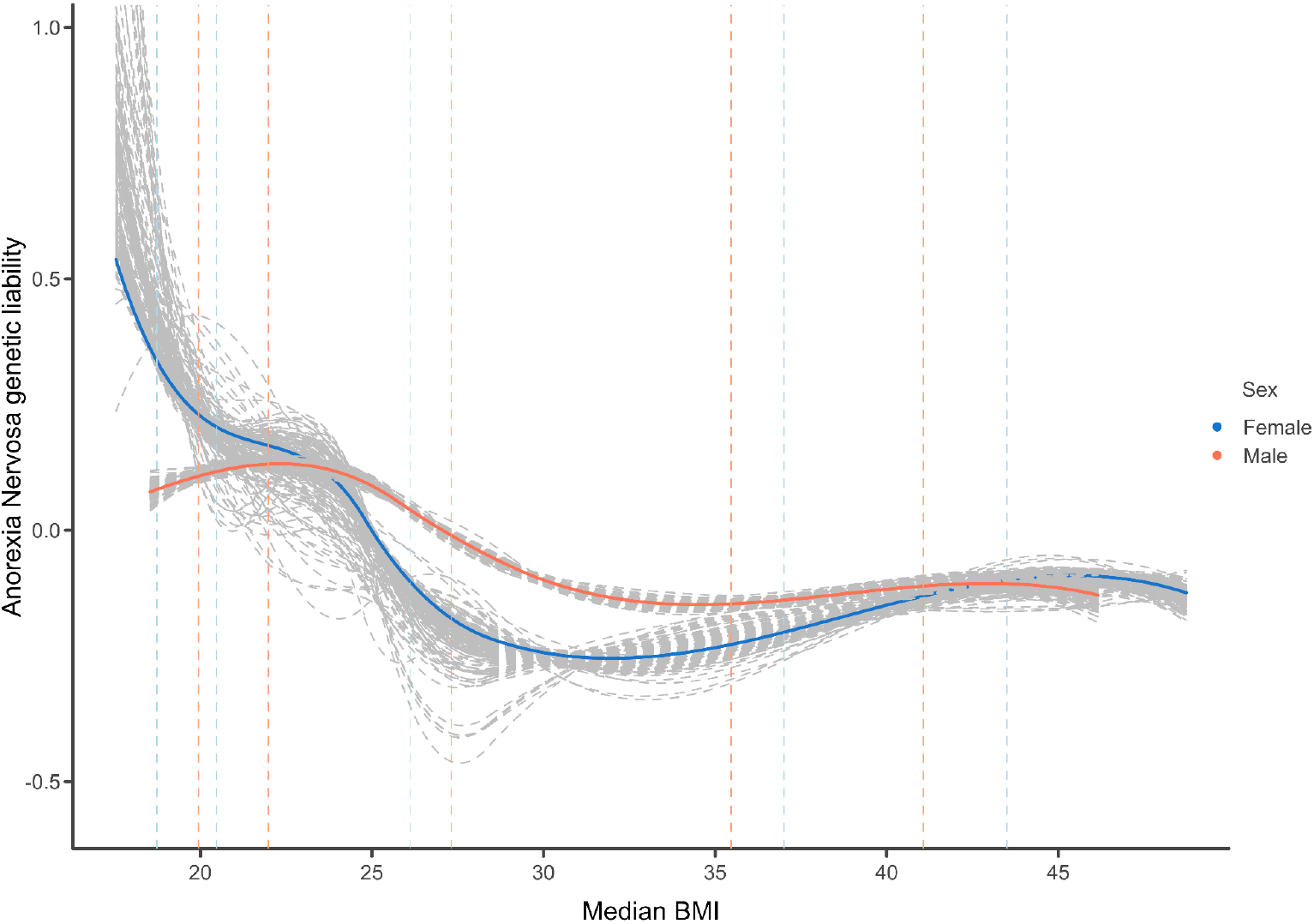
Sex-stratified genetic relationship between BMI and anorexia nervosa ***Note***: *median BMI per bin is plotted on the x-axis and the y-axis represents the expected liability of anorexia nervosa given BMI on a relative scale (scale values cannot be compared between outcomes). Dashed lines represent BMI values at the 1st, 5th, 50th, 95th and 99th percentile of the data for each sex. Results are based on the cubic spline estimator*

### Sleep duration

Low and non-significant genetic correlation estimates around −0.1 (SE = 0.04-0.06)^18,27^ have been reported between sleep duration and depression, and around ±0.1 (SE = 0.03-0.05)^18,27^ between sleep duration and ADHD, but our analyses suggest a clear nonlinear U-shaped relationship. The expected value of *a*_*dep*/*adhd*_ (*E*[*a*_*dep*/*adhd*_]) is a nonlinear function of *a*_*bim*_ (*E*[*a*_*bim*_]). In the case of anorexia, genetic correlations with sleep duration are also low and non-significant (r_g_ = <-0.1, SE = 0.04-0.06)^18,27^ though our analyses suggest no underlying nonlinear relationship (Figure 4). Full correlation results are reported in Supplementary Figures 9-11 and Tables 7-9).

**Figure 4:**
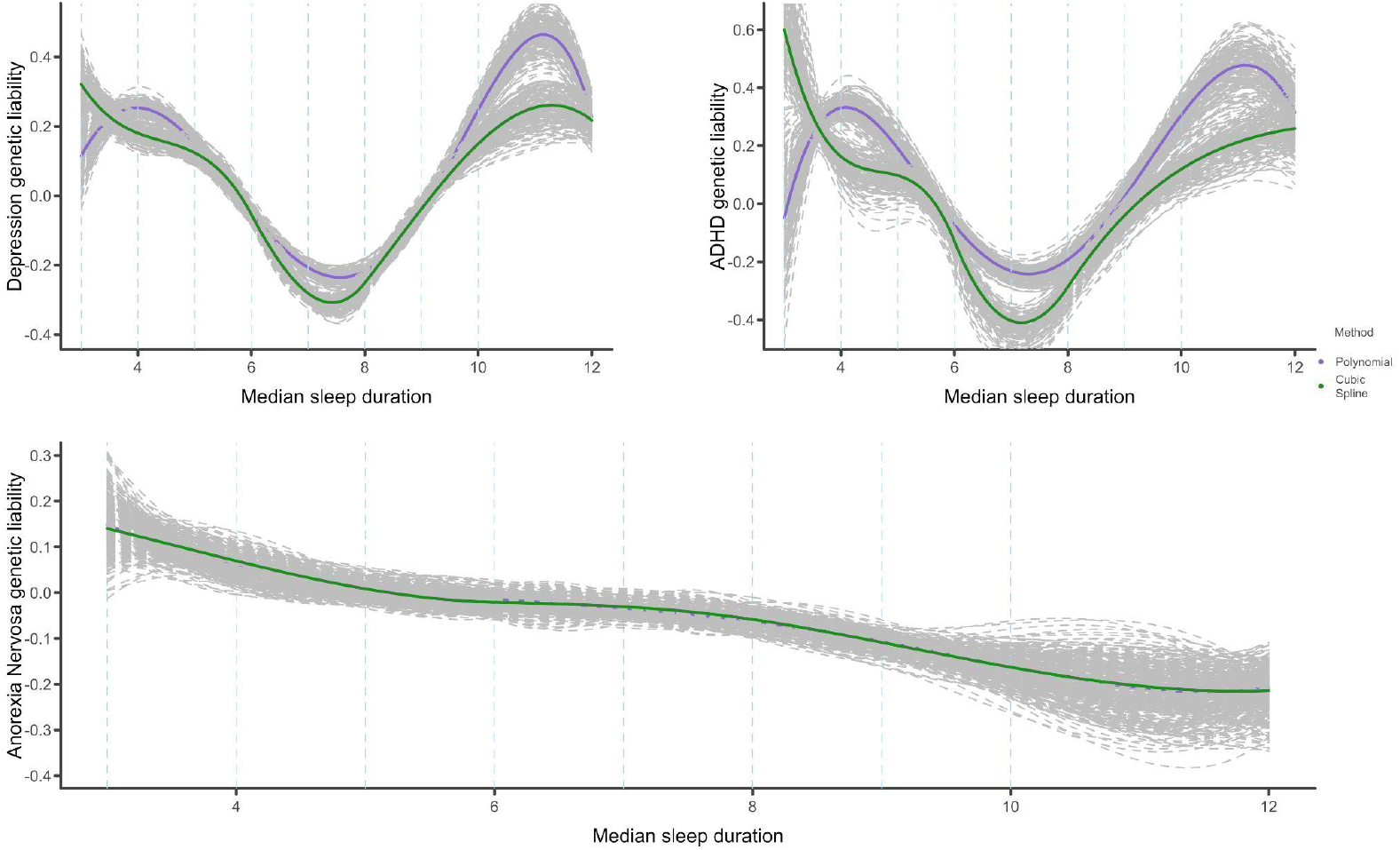
Genetic relationship between sleep duration and psychiatric disorders ***Note***: *median sleep duration per bin is plotted on the x-axis and the y-axis represents the expected liability of the relevant disorder given sleep duration on a relative scale (scale values cannot be compared between outcomes). Dashed lines represent maximum sleep duration values at each of the 9 bins*

### Height

No nonlinear relationships were observed between height and any of the psychiatric disorders (Figure 5). Aggregate estimates suggest no significant genetic correlations between height and depression (r_g_ = −0.05, SE = 0.02, P = 0.02) ^26^, ADHD (r_g_ = −0.065, SE = 0.03, P = 0.02), and anorexia (r_g_ = 0.00, SE = 0.03, P = 0.99) ^24^. The relationship between height and ADHD and height and depression appear linear, and entirely absent in the case of anorexia. The flat pattern observed with anorexia is the result of a 0 genetic correlation with height, while ADHD and depression have very weak negative genetic correlations with height. Full case-control correlation results are reported in Supplementary Figures 12-14 and Tables 10-12).

**Figure 5:**
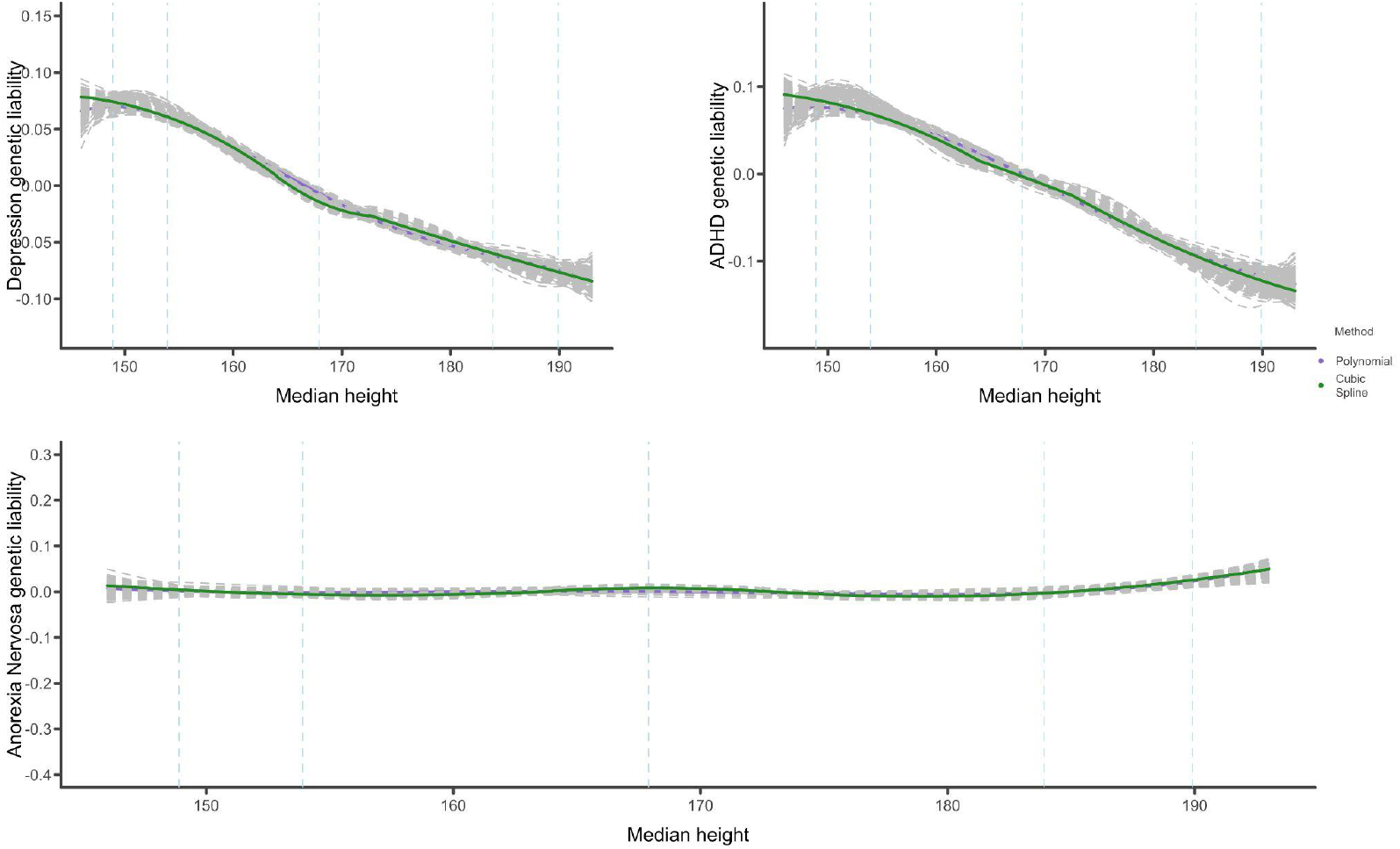
Genetic relationship between height and psychiatric disorders ***Note***: *median height per bin is plotted on the x-axis and the y-axis represents the expected value of the relevant disorder given height on a relative scale (scale values cannot be compared between outcomes). Dashed lines represent height values at the 1st, 5th, 50th, 95th and 99th percentile of the data*

## DISCUSSION

In the current study, we introduce and apply a novel method to estimate nonlinear genetic relationships between continuous and binary traits using GWAS summary statistics. The approach leverages genetic correlations between different cross sections of a continuous trait (e.g., BMI or sleep duration) and a binary outcome (e.g., depression, ADHD, anorexia nervosa) to infer the shape of their genetic relationship. This methodology departs from traditional approaches like genetic correlations which only detect the linear component of a relationship, offering a more nuanced understanding of the underlying genetic relationship between traits. Our method does require specialised additional GWAS of the continuous trait in question, but then subsequently only relies on GWAS summary data to address the limitations of standard genetic correlation estimates, which, while informative about the overall genetic relationship between traits, fail to capture the potential complexity of these relationships. By binning the continuous trait into multiple quantiles and estimating genetic correlations between each pair of bins and the binary trait, we can infer the functional relationship between the genetic liabilities of the two traits across the distribution of the continuous trait. This approach allows us to detect and model nonlinearities that might otherwise remain obscured in aggregate analyses.

It is important to note that this method is purely observational and does not seek to answer questions about how or why the nonlinearities exist; rather, it seeks to establish whether or not they do. Further questions about why they exist constitute causal questions which we consider to be beyond the scope of the current study. Crucially, the presence of a detectable nonlinear genetic relationship does not suggest a nonlinear causal relationship exists between traits. In fact it might suggest that variables like BMI aren’t coherent traits at all, with low and high BMI having their own set of distinct causes, consequences and relationships. Prior controversy around nonlinear mendelian randomization estimators suggests nonlinear causal inferences based on genetic data may be fraught ^28^. Finally, phenotype definitions may impose constraints on the relationship between traits, and thus impact findings. For example, symptoms of depression include weight gain/loss, and insomnia/hypersomnia, which may have influenced our findings. As such, the resulting shapes can be an artefact of the trait definitions. Importantly, our use/application of genetic data allows us to investigate this in traits that have not been measured in the same individuals.

### Application and Findings

When applied to the genetic relationship between BMI and psychiatric disorders (depression, ADHD, anorexia nervosa), our method revealed nonlinearities that traditional genetic correlation approaches might miss. For example, the nonlinear relationship between BMI and depression, characterised by a J-shaped curve, suggests that different genetic factors may influence depression risk at different points along the BMI phenotypic spectrum. We observe similar relationships in the association between self-reported sleep duration and ADHD and depression. This does not appear to generalise to anorexia as we observe no underlying nonlinearity in the relationship with sleep duration. These findings are in line with results that showed that while long and short sleep duration were genetically correlated with depression, total sleep duration was not ^18^, as well as studies that have used twin studies to show that the heritability of BMI varies across its phenotypic range ^29^. The results highlight the potential for differential genetic expression or the involvement of distinct genetic variants at the extremes of the BMI and sleep duration distributions. Analysis of the relationship between BMI and anorexia nervosa uncovered a more complex pattern than previously reported. While traditional analyses suggest a negative linear correlation, our analyses shows that this relationship is primarily confined to lower BMI ranges, with a plateau or even reversal in higher BMI ranges, particularly in females. This insight could have significant implications for understanding the genetic aetiology of anorexia nervosa, especially concerning the role of body weight in modulating genetic risk. The robustness of our approach was further demonstrated by sensitivity analysis that reveal the relationship between height and psychopathology to be entirely linear or absent.

As discussed in the introduction, in genetics, non-linearity is frequently invoked to describe the relationship between allele and outcome. Our methods and findings imply that genetic effects on traits may also operate in a manner that depends on the level of the trait. In statistical genetics these effects are seen as gene-environment interaction and studied using “reaction norm models”. So while we detect non-linearities, these could be viewed as part of underlying processes such as genotype-covariate correlation and interactions (GCCI) and residual-covariate correlation and interaction effects (RCCI), which can be estimated by whole-genome reaction norm models ^30^. However, we do not investigate genotype-environment interplay (GxE) in the typical sense as we consider no external environment, rather we split individuals along the outcome variable itself. Future work should investigate how the presence of gene-environment interactions, particularly those detected in reaction norm models, and non-linear relationships between traits are simply two ways of detecting a single phenomenon or whether they are separate processes that can occur independently or concurrently.

### Limitations and considerations

An important limitation is that nonlinear dependencies are not symmetric; whether or not the mean of *x* depends nonlinearly on *y*, it does not imply that the mean of *y* depends nonlinearly on *x*. In our application we stratify BMI, and indirectly the BMI liability, but we do not stratify the depression or ADHD or anorexia liability. There is therefore a risk of overgeneralizing a finding that “the expected value of *y* (*E*[*y*]) is, or is not, a nonlinear function of *x* (*f*(*x*))” to “there does, or does not exist, a nonlinear relationship between *x* and *y*”.

Secondly, detection of nonlinearity is dependent on genetic heterogeneity in the trait that is binned/split. If all genetic effects on that trait are truly additive and homogeneous across the trait spectrum, any nonlinear effects it has on a second trait becomes undetectable. This is problematic if the second trait is binary; however if the second trait is continuous it could be stratified instead and nonlinearity may still be estimated. Our method will not work on two traits that are fully additive/homogeneous, though for the genetic influences on the trait to be fully additive requires that there are no non-linear influences from other heritable traits. Thus it is exceedingly unlikely two fully additive traits are non-linearly related (this would require non-linear effects to systematically be canceled out).

Thirdly, careful consideration of specified parameters must be undertaken as this can influence the resulting shape of a relationship. For example the choice of nonlinear plotting function (e.g. cubic-spline vs polynomial), or indeed the specification of y-axis limits, can influence the perception of the resulting shape produced. Additionally, they are scale dependent; the choice of specific mapping from discrete bin, to numerical variable (e.g. bin number vs median BMI for the bin) and its resulting distribution on the x-axis will influence the shape of the estimated nonlinear function. For example, compared to plotting bin numbers which are equidistant to adjacent bins, plotting the median BMI value in each bin affects the distance between subsequent points especially at the right tail of the distribution where they become much larger, resulting in a different estimated function. The reduced power from smaller samples in the tail bins also introduces local uncertainty in the tails of the nonlinear relationship.

Fourthly, for some scales (like hours slept) an obvious natural scale might exist, but for other scales there may be no natural way to map discrete bins to a numeric scale. Changing the scale of one of the variables will change the shape of the function. Especially in the tails, if the function overfits the data the shape of the relation could become dependent on scale. We recommend using a natural scale if available, and if not a careful consideration across a number of competing reasonable scales.

Finally, the choice of the number of bins as well as how they are specified are fairly arbitrary and dependent on the continuous phenotype of interest, and how many bins it can be stratified into while preserving enough power in each for the GWASs. Care should be taken in the decisions, and interpretation of any resulting shape underlying the relationship between traits.

### Implications and Future Directions

The ability to model and detect nonlinear genetic relationships has several important implications. First, it challenges the prevailing assumption of linearity in genetic epidemiology, suggesting that more complex models may be necessary to fully capture the nature of genetic associations. Second, the detection of nonlinear relationships can inform the development of more targeted interventions, as different segments of the population may be genetically predisposed to different risks.

The method can be applied to other traits and conditions, offering a new tool for exploring genetic complexity in various domains of biomedical research. Future work could involve refining the binning strategy, exploring alternative functional forms beyond polynomials, applying this method to larger and more diverse datasets, extensions into longitudinal modelling, as well as further investigations into the biological mechanisms underlying these associations.

In conclusion, our study presents a novel methodological approach that is a point of departure for the exploration of nonlinear genetic relationships between traits. By moving beyond the assumption of linearity, we have uncovered new insights into the genetic architecture of complex traits, with potential implications for both research and clinical practice. This method represents a significant step forward in the field of genetic epidemiology, providing a framework for more accurately modelling the genetic underpinnings of human traits. Theories for the existence of nonlinearity include definitions of psychiatric nosology, while a genetic reason could be that this is a form of GxE or reaction norm; future work untangling the processes underlying nonlinearities are crucial. We do however show that they are detectable.

## ACKNOWLEDGEMENTS

WAA is supported by the National Institute Of Mental Health of the National Institutes of Health under Award Number R01MH120219 and a Dutch Ministry of Education, Culture and Sciences (OCW) Talent and Early Development Grant. MGN is supported by the National Institute Of Mental Health of the National Institutes of Health under Award Number R01MH120219.

