## Supplementary Text for "Detecting Non-linear Dependence through Genome Wide Analysis"

**Simulations**

We used GWASbrewer [1] to simulate 19490 variants with an LD structure that matches chromosome 19, for 60.000 individuals, using the LD structure provided in the GWAS Brewer package.

Based on the known LD structure we compute LD scores (the sum of squared pairwise correlations with the focal variant). We sample effect sizes from a multivariate normal distribution for 2 independent traits and simulate environmental effects (fixing SNP heritability at 25%). We then introduce a (nonlinear) relationship between traits to validate our methods.

For two traits y1 and y2 we introduce the following relationships: **curve up (a U shape)**, where for half of simulated individuals y1 increases as y2 decreases, while for other simulated individuals y1 increases as y2 increases; **linear-positive** where y2 has a positive linear relationship with y1; **linear-negative** where y2 has a negative linear relationship with y1; **flat-then-up** where y2 and y1 are unrelated when y1 is negative, but positively linearly related when y1 is positive; **down-then-flat** where y2 and y1 are unrelated when y1 is negative but negatively linearly related when y1 is positive; and finally **uncorrelated**, where y2 and y1 are uncorrelated.

We bin y2 with bins bounded at the 100th, 90th, 80th, 70th, 60th, 40th, 30th, 30th, 20th, 10th and 0th percentile, perform bin vs bin GWASs (abbreviated to a simple correlation as there is no confounding), and then compute genetic correlations between these GWASs and the GWAS of y1 by performing abbreviated LDscore regression. We then applied cor2curve, the function that implements our nonlinear estimator in TriGenometry, using a polynomial function. Results are presented in Supplementary Figure 1. Red and blue dots represent the observed genetic liability of y1 in each bin of the observed variable y2, while the black line represents the estimated (non)-linear relationship. Note that due to relative scaling and the fact that we perform bin-wise GWAS of the phenotype rather than of the genetic liability, we expect to only retrieve the shape of the relationship, not the absolute values of the liability of y1 given a value of y2.

For practical computational reasons our simulation uses a regularised, somewhat simplified LD structure, a small genome, and simplified GWAS and LDSC methods. Our simulation confirms that the presence of a nonlinear phenotypic relationship, where y1 influences the conditional genetic liabilities of y2 in a nonlinear manner, can be detected. Our simulation does not ensure this is the only process that gives rise to an apparent nonlinear relationship; patterns of gene-environment interaction and other unmodeled processes may introduce apparent nonlinear relations. Our simulation also does not ensure that all nonlinear relations are detected. For example we model y2 conditional on y1, if the true relationship is reversed and y2 has a non-linear effect on y1, there is no guarantee the nonlinearity would be detected. Given the number of possible nonlinear relationships between traits is large, creating a “garden of forking paths”, we advocate people perform their own domain specific simulations, incorporating substantive knowledge of potential sources of potential bias.

**Supplementary Figure 1:** Estimated nonlinear genetic relationships (black line) and phenotypic expectations of y1, given y2 ( phenotypic ground truth).

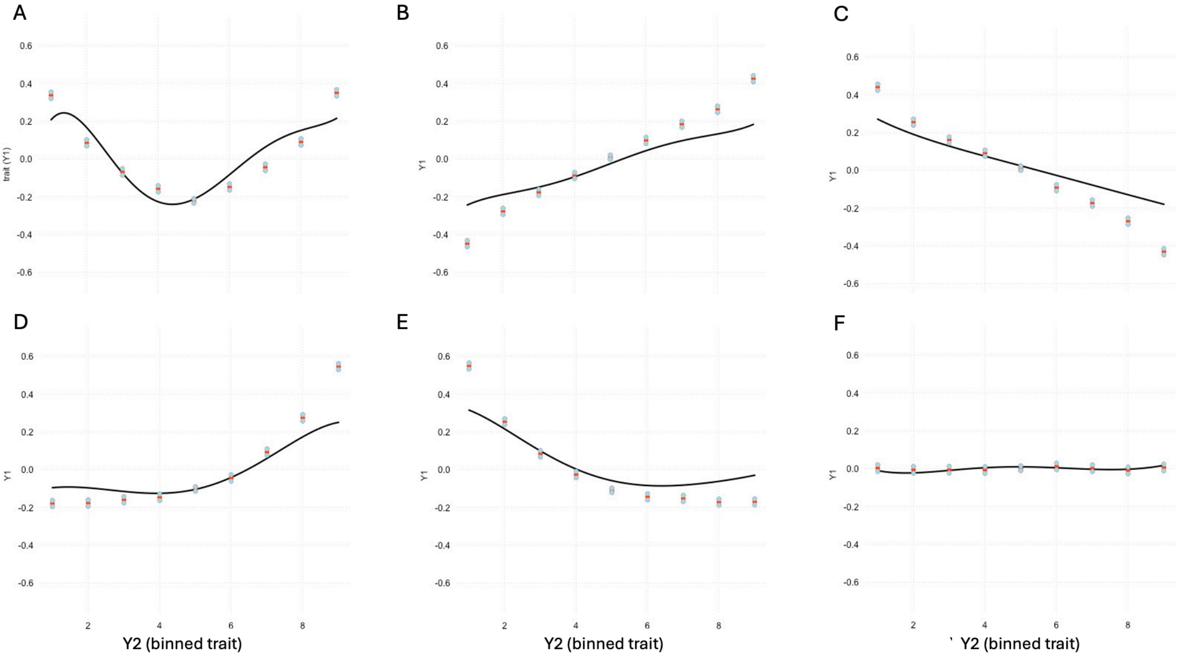


**Simulated violation of assumptions**

We outline various assumptions, one of these states that in order to detect a nonlinear relationship, genetic influences on the variable that is binned ($y2$ in our notation) must have detectable heterogeneity across levels of $y2$. Here we simulate data where this assumption is violated, in order to illustrate how a false negative may arise.

We simulate $y2$ as:

$$y2 = \sum_{1}^{m}(\beta_m * snp_m) + e$$

Where β are sampled from a normal distribution and $snps$ $1$ to $m$ are simulated single nucleotide polymorphisms.

And we simulate y1 as -2*$y2$, when $y2$ goes up over 0.5 (a “flat then down” pattern).

$$y1 = -2y2 * (y2 > .5)$$

We then perform segmented GWAS of $y2$, and use LDscore regression to correlate those to a GWAS of $y1$. As outlined in the assumptions section of the methods, the type of relationship simulated here is a type of nonlinear relationship our method fails to detect (**Supplementary Figure 2**), because the SNP effects on y2 are fully additive.

**Supplementary Figure 2**: when assumptions about the genetic architecture of the segmented trait are violated the observed nonlinear relationship mean $y1$ values for bins of $y2$ goes undetected.

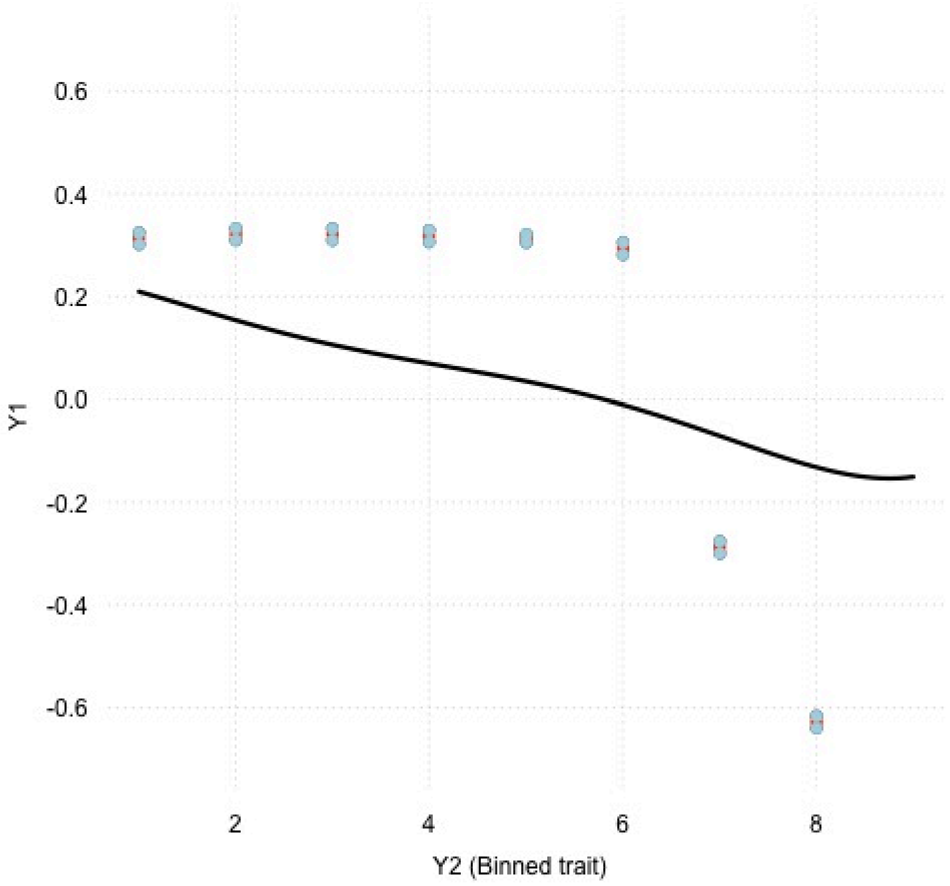


The asymmetry illustrated here does not imply our test somehow uncovers directionality in the relationship between $y2$ and $y1$, as either confounders with a nonlinear effect on $y2$ and $y1$, or any other sources of nonlinear genetic effects on $x$ are detected by our method.

**Supplementary Figure 3**: correlation between BMI and depression

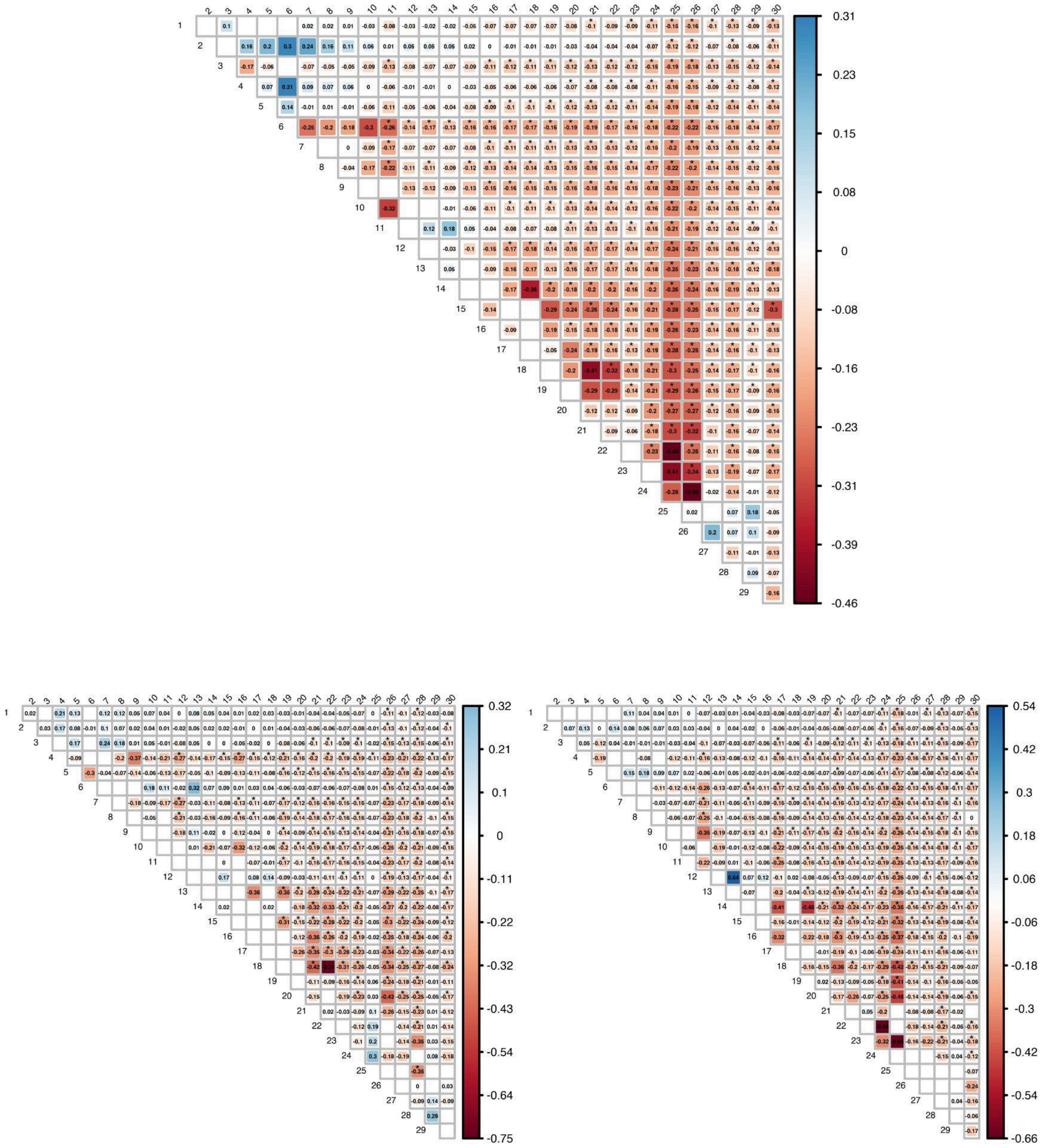


Note: Diagonal labels represent BMI GWAS case bin numbers while top labels represent control bins. Blank cells are where correlation estimates with depression could not be estimated. Significant (at p< 0.05) estimates are indicated with * above. Top plot is combined sample, bottom left is male, and bottom right is female

**Supplementary Figure 4**: correlation between BMI and ADHD

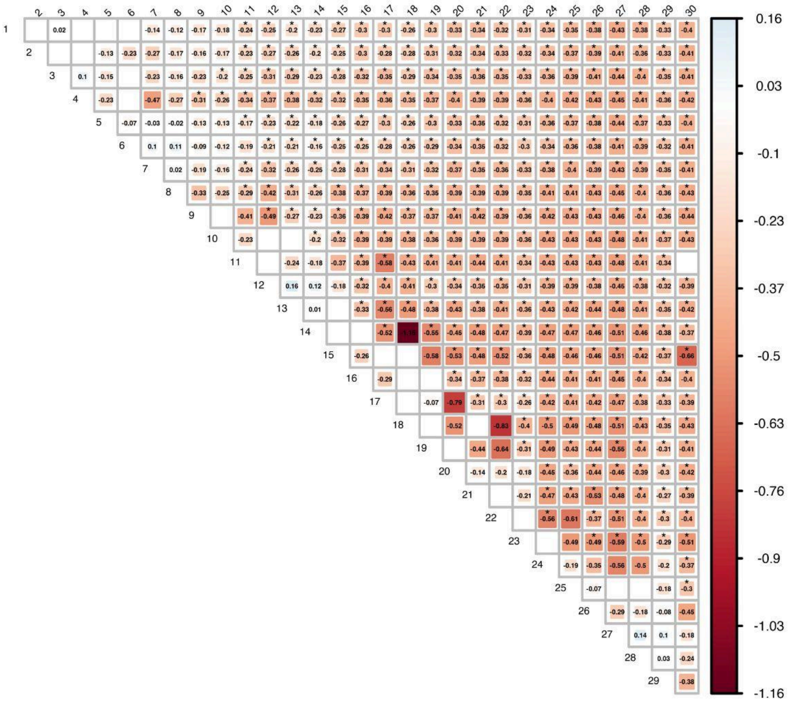

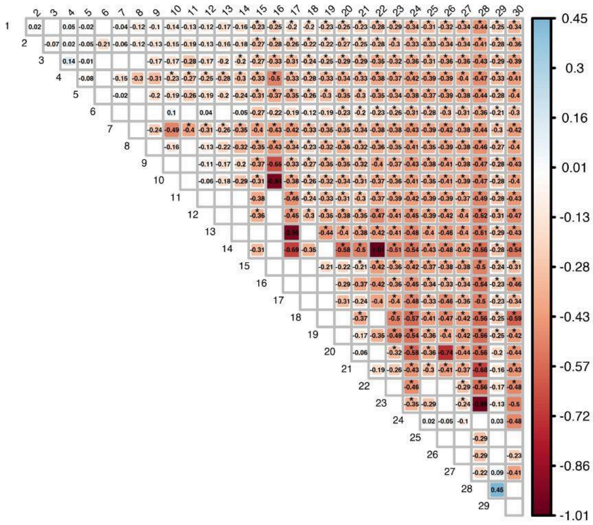

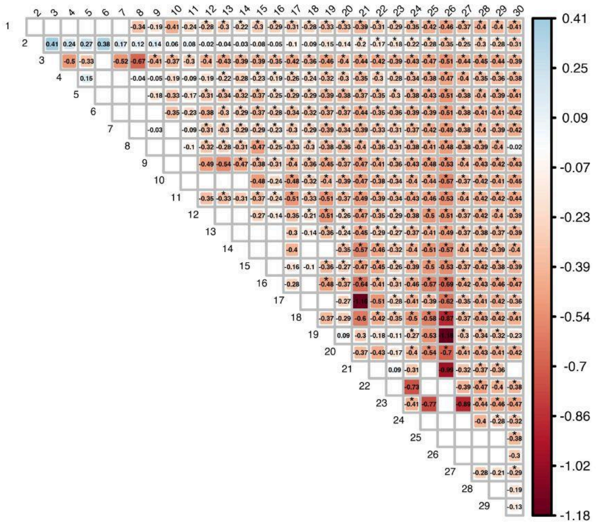

Note: Diagonal labels represent BMI GWAS case bin numbers while top labels represent control bins. Blank cells are where correlation estimates with ADHD could not be estimated. Significant (at p< 0.05) estimates are indicated with * above. Top plot is combined sample, bottom left is male, and bottom right is female

**Supplementary Figure 5**: correlation between BMI and anorexia nervosa

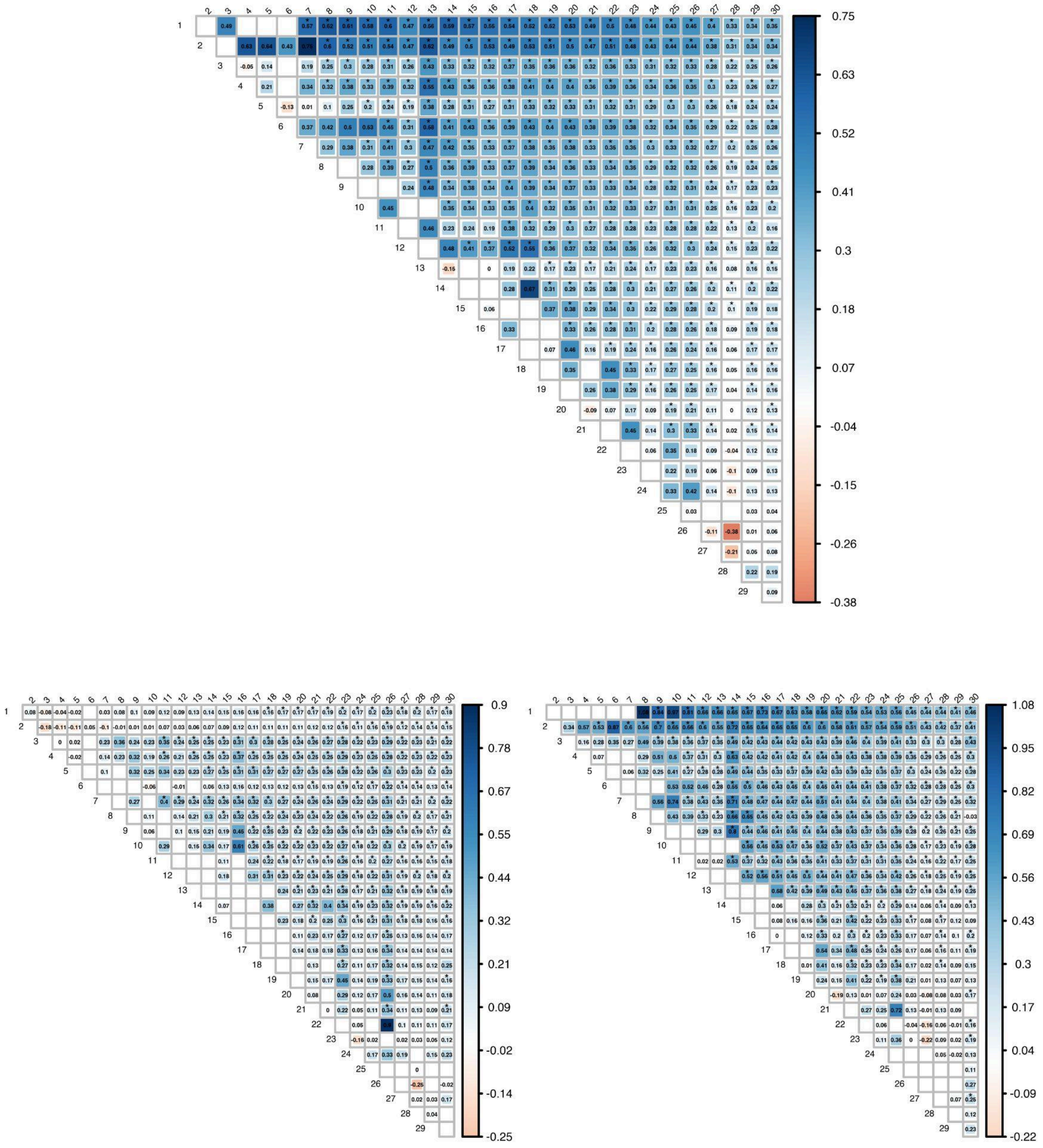

Note: Diagonal labels represent BMI GWAS case bin numbers while top labels represent control bins. Blank cells are where correlation estimates with anorexia could not be estimated. Significant (at p< 0.05) estimates are indicated with * above. Top plot is combined sample, bottom left is male, and bottom right is female

**Supplementary Figure 6:** sex-stratified non-linear association between BMI and anorexia

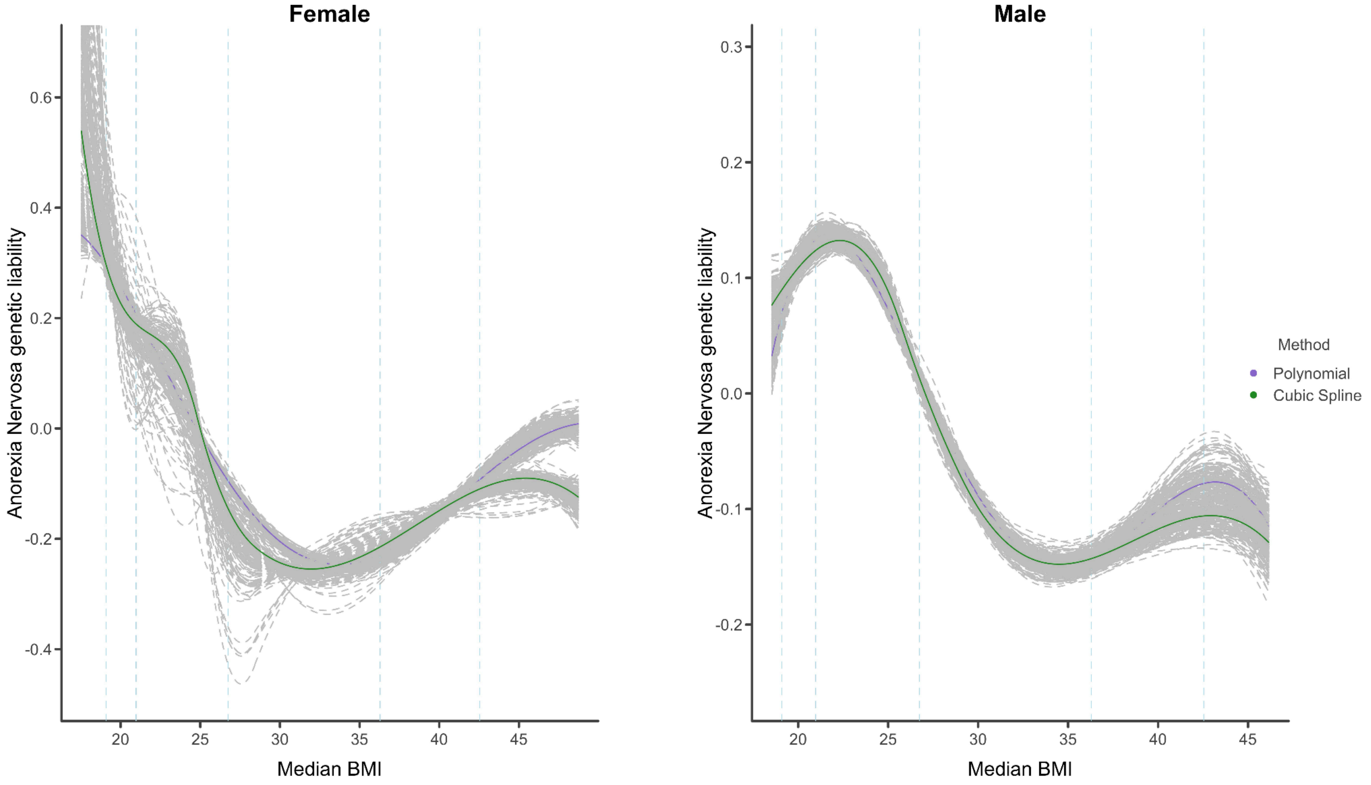


**Supplementary Figure 7:** sex-stratfied results of non-linear association between BMI and ADHD

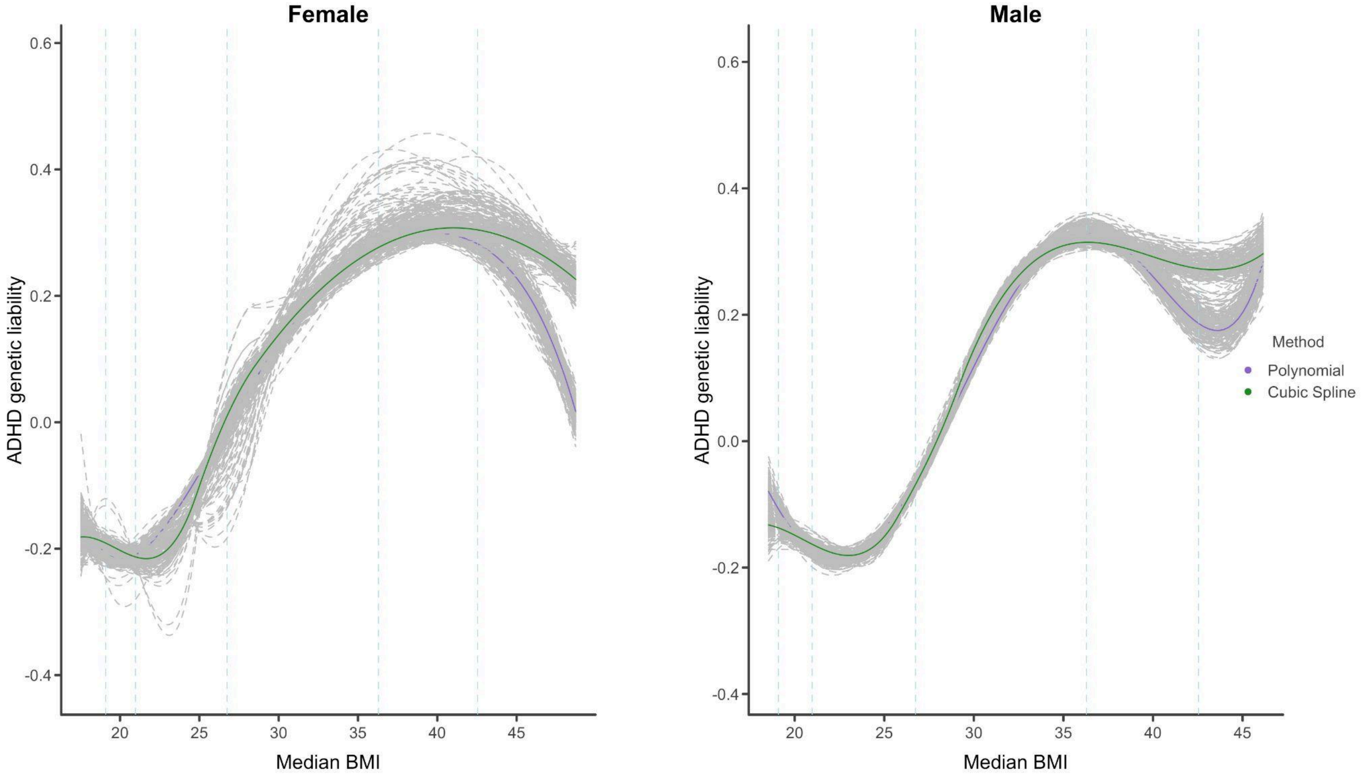

**Supplementary Figure 8:** sex-stratfied results of non-linear association between BMI and depression

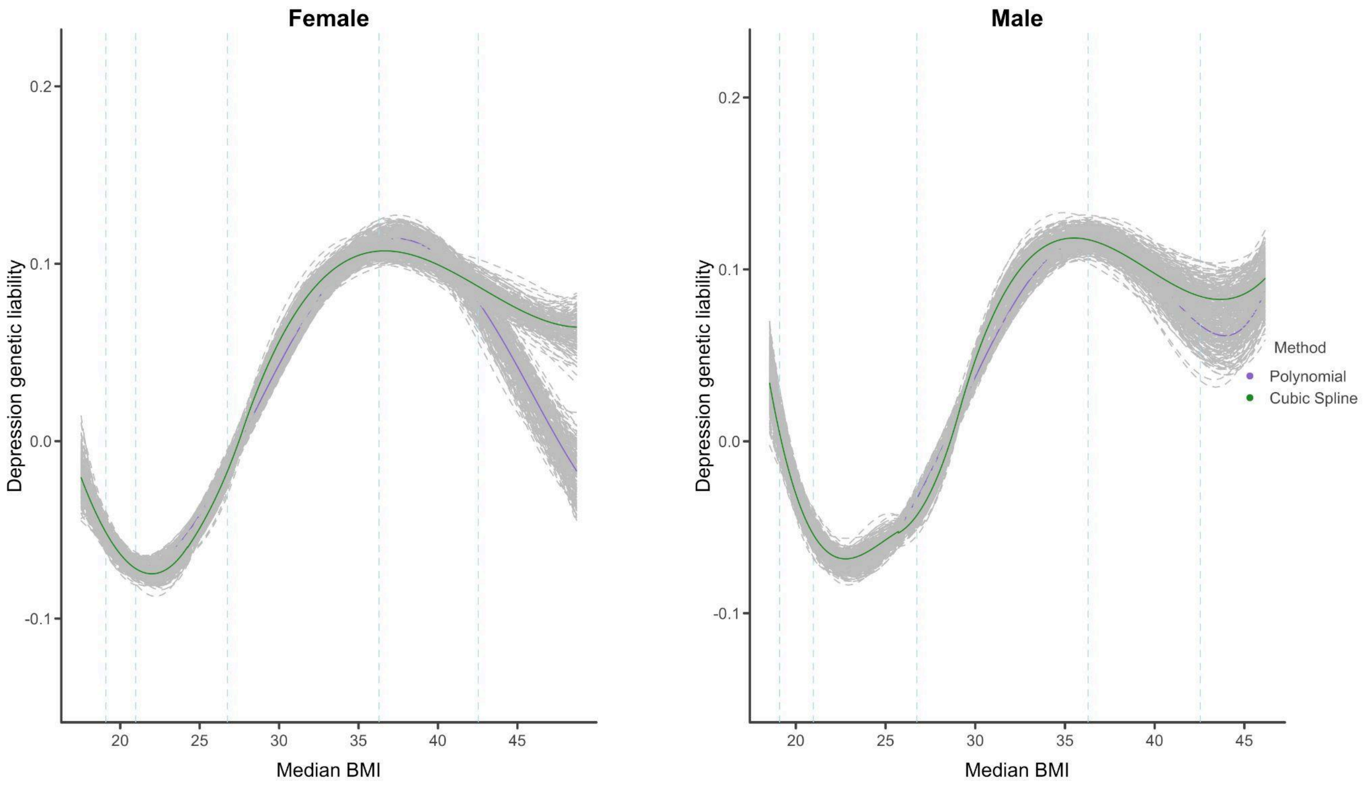


**Supplementary Figure 9**: correlation between sleep and depression

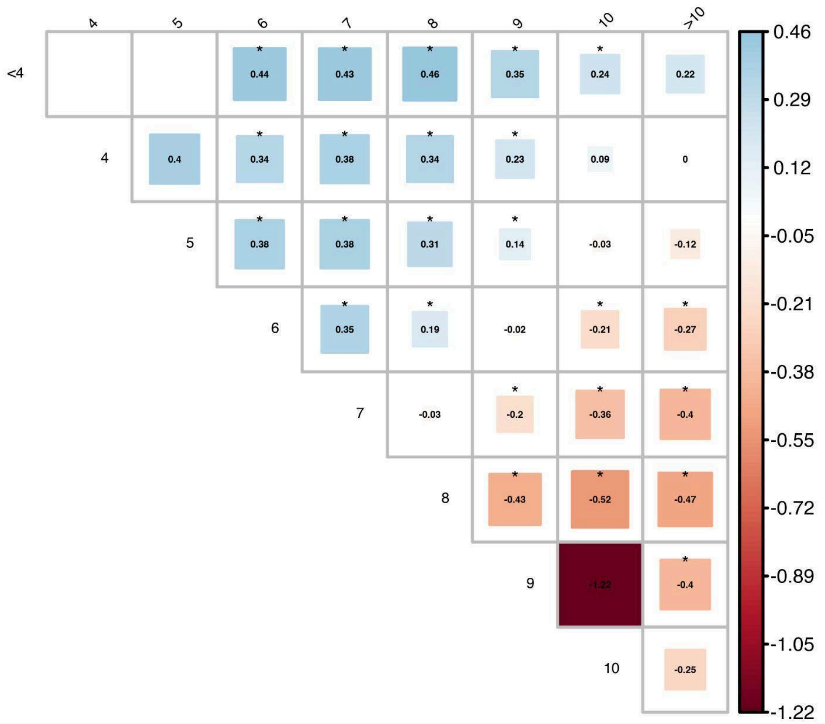


Note: Diagonal labels represent number of hours slept for cases, while top labels indicate number of hours slept for controls.Blank cells are where correlation estimates with depression could not be estimated. Significant (at p< 0.05) estimates are indicated with * above

**Supplementary Figure 10**: correlation between sleep and ADHD

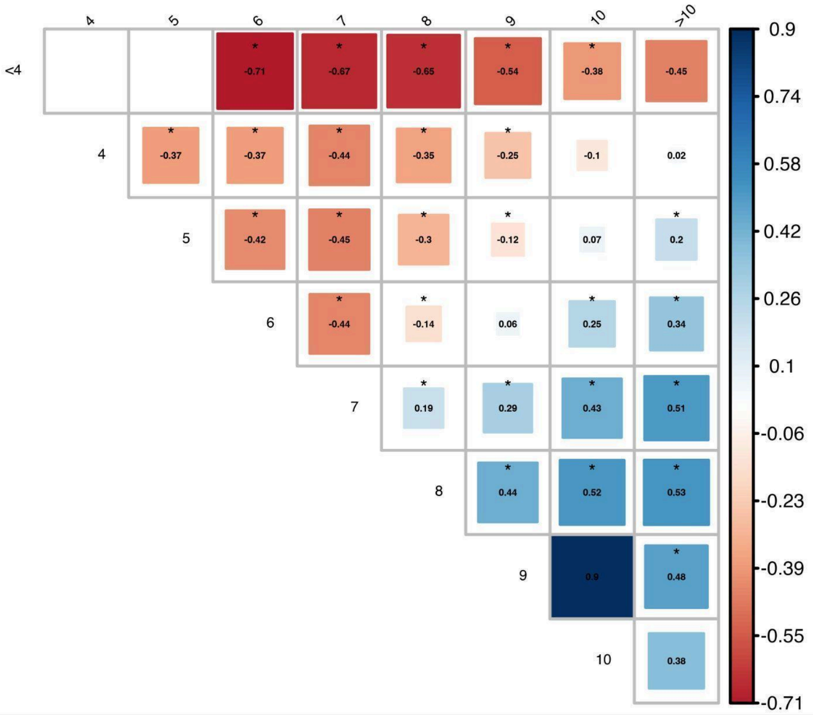


Note: Diagonal labels represent number of hours slept for cases, while top labels indicate number of hours slept for controls.Blank cells are where correlation estimates with ADHD could not be estimated. Significant (at p< 0.05) estimates are indicated with * above

**Supplementary Figure 11**: correlation between sleep and anorexia

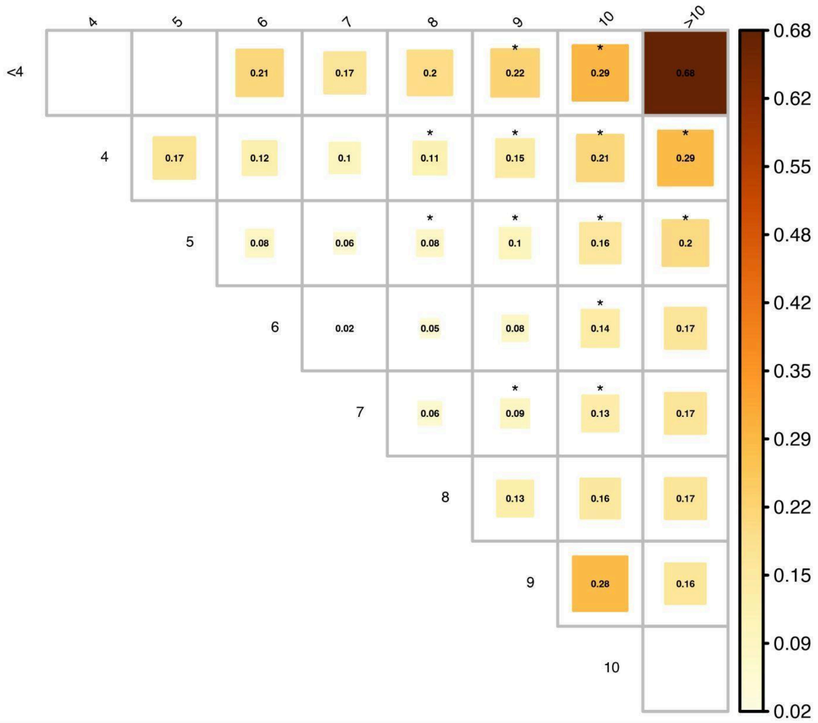


Note: Diagonal labels represent number of hours slept for cases, while top labels indicate number of hours slept for controls. Blank cells are where correlation estimates with anorexia could not be estimated. Significant (at p< 0.05) estimates are indicated with * above

**Supplementary Figure 12**: correlation between height and depression

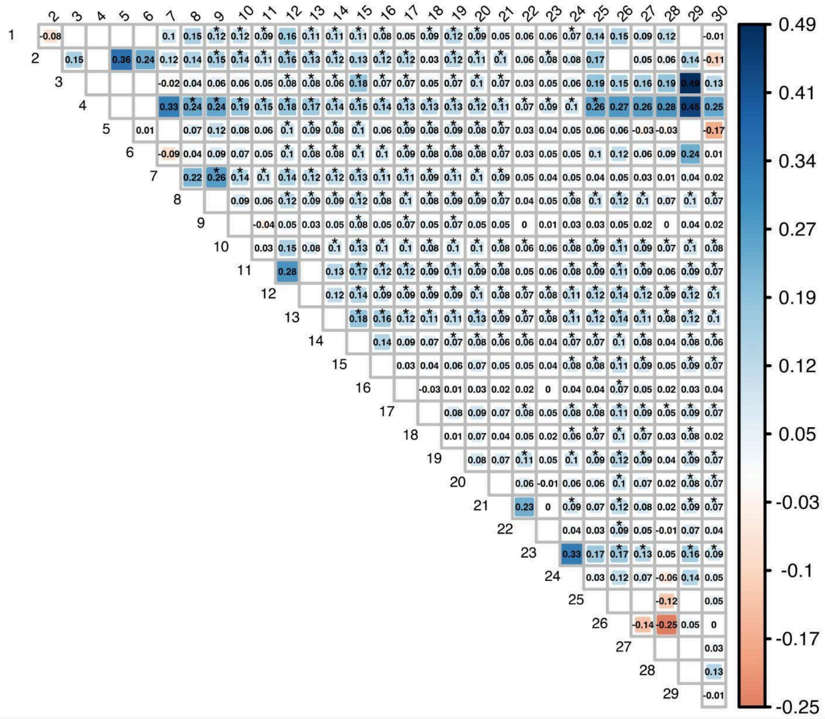


Note: Diagonal labels represent height GWAS case bin numbers while top labels represent control bins. Blank cells are where correlation estimates with depression could not be estimated. Significant (at p< 0.05) estimates are indicated with * above

**Supplementary Figure 13**: correlation between height and ADHD

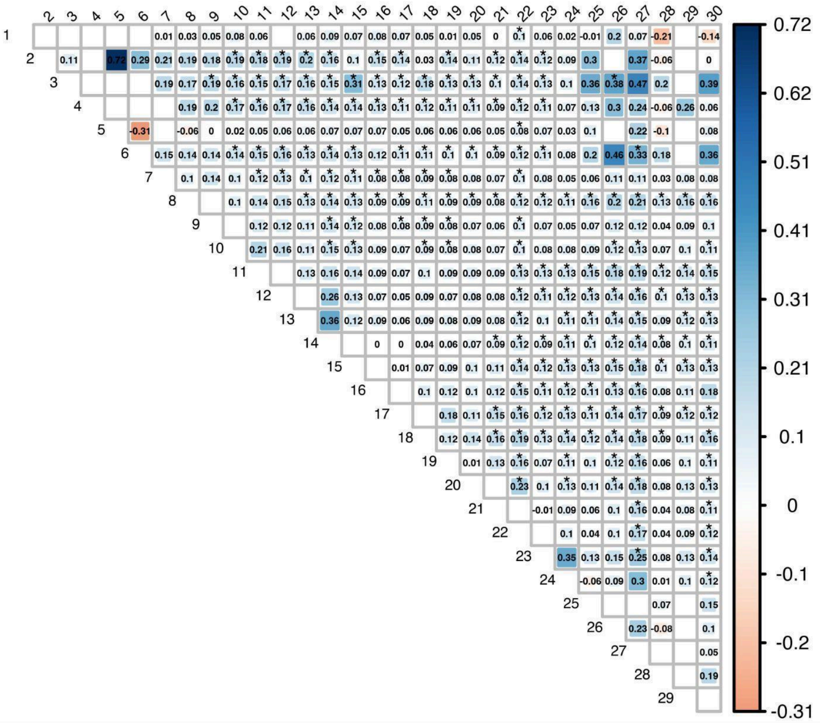

Note: Diagonal labels represent height GWAS case bin numbers while top labels represent control bins. Blank cells are where correlation estimates with ADHD could not be estimated. Significant (at p< 0.05) estimates are indicated with * above

**Supplementary Figure 14**: correlation between height and anorexia

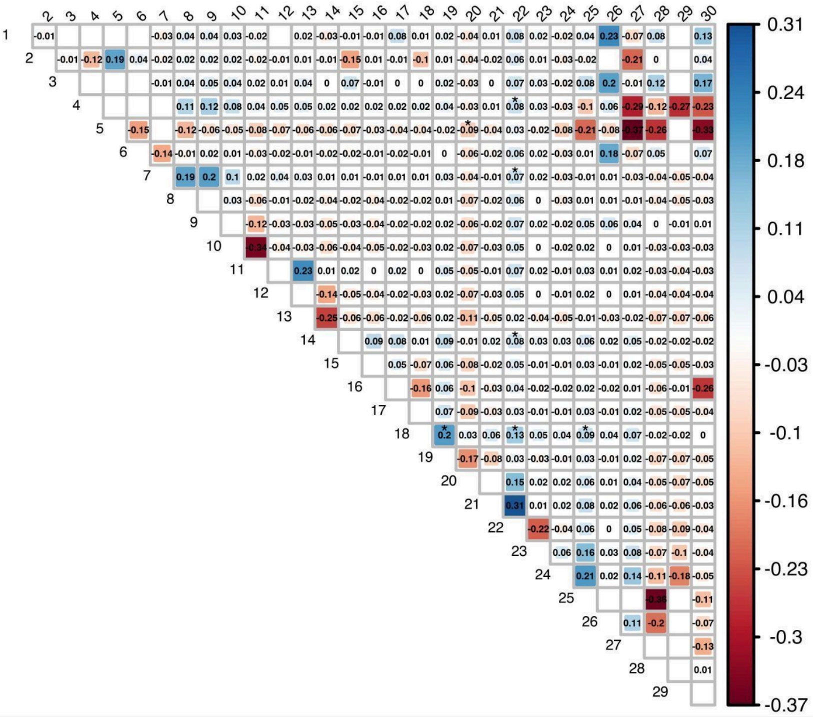

Note: Diagonal labels represent height GWAS case bin numbers while top labels represent control bins. Blank cells are where correlation estimates with anorexia could not be estimated. Significant (at p< 0.05) estimates are indicated with * above

**REFERENCES**

1. Morrison J. GWASBrewer: An R Package for Simulating Realistic GWAS Summary Statistics. *Genet Epidemiol*. n/a(n/a). doi:10.1002/gepi.22594
